## Supplementary figure 1 for "The cardiac, respiratory, and gastric rhythms independently modulate motor corticospinal excitability in humans"

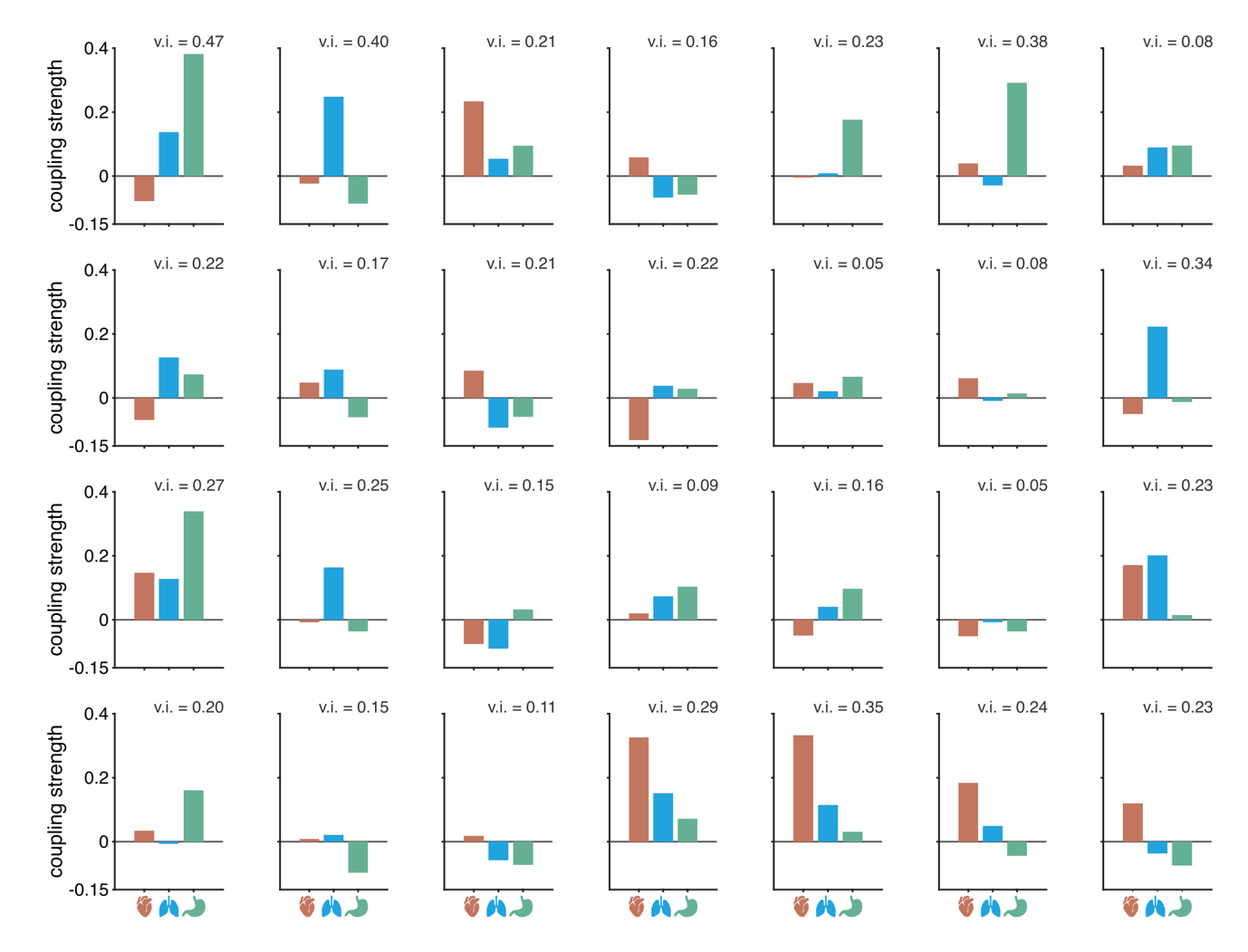

**Supplementary figure 1.** Individual interoceptive profiles of all 28 participants. For each participant the coupling strength between each of the visceral rhythms (red = cardiac, blue = respiratory, green = gastric) and the amplitude of the MEP is plotted. The variability index (v.i.) of each participant reflects how homogeneous or heterogenous coupling strength was across the different organs, with larger values indicating more heterogeneity. Interoceptive profiles show large variability, with both homogenous (e.g. coupling to all, or to none, of the visceral rhythms) and heterogenous (coupling to one or two, but not all visceral rhythms) profiles.
